# A polyclonal allelic expression assay for detecting regulatory effects of transcript variants

**DOI:** 10.1101/794081

**Authors:** Margot Brandt, Alper Gokden, Marcello Ziosi, Tuuli Lappalainen

## Abstract

We present an assay to experimentally test regulatory effects of genetic variants within transcripts using CRISPR/Cas9 followed by targeted sequencing. We applied the assay to 35 premature stop-gained variants across the genome and in two Mendelian disease genes, 33 putative causal variants of eQTLs and 65 control variants. We detected significant effects generally in the expected direction, demonstrating the ability of the assay to capture regulatory effects of eQTL variants and nonsense-mediated decay triggered by premature stop-gained variants. The results suggest a utility for validating transcript-level effects of genetic variants.

## Background

The interpretation of functional effects of common and rare variants in the human population is a major objective in human genetics and genomics. Despite the success of mapping genetic associations to complex traits by genome-wide association studies (GWAS) and interpreting their effects on gene expression by expression quantitative trait loci (eQTL) studies (Battle et al., 2014; Grundberg et al., 2012; GTEx Consortium et al., 2017; Lappalainen et al., 2013; Nicolae et al., 2010), the causal variants at GWAS loci and eQTLs are usually unknown due to linkage disequilibrium (LD). Statistical fine-mapping methods (Hormozdiari et al., 2014; Wellcome Trust Case Control Consortium et al., 2012; Wen et al., 2015) can help narrow down causal variants, but experimental validation of the performance of these methods is lacking. For rare variants, functional interpretation has distinct challenges even in the well-annotated coding regions. Rare disease studies often result in 100s to 1000s of potential disease-causing variants identified from whole exome sequencing, and prioritization based on their functional effect is essential for research and clinical use (Gilissen et al., 2012).

Thus, there is a need for experimental methods to confirm the effects of common and rare variants. Methods such as massively parallel reporter assays (MPRAs) (Tewhey et al., 2016; van Arensbergen et al., 2019), which couple regulatory sequences with an expression-correlated reporter, are high-throughput approaches for finding active regulatory variants outside of the gene body, and analogous methods exist for variants affecting splicing (Ke et al., 2011; Wang et al., 2012). However, the results of the assays show low concordance with eQTL data (Tewhey et al., 2016; van Arensbergen et al., 2019), perhaps due to taking the variant out of its genomic context. Furthermore, MPRAs are not suited to testing variants within the transcript that can affect gene expression levels via post-transcriptional mechanisms, e.g. RNA stability. Regulatory variants are strongly enriched not only in promoters and enhancers, but also in UTRs and other transcript annotations (Lappalainen et al., 2013), emphasizing the need for a method to validate them. Similar mechanisms also apply to a subset of rare disease variants, where stop-gained and frameshift variants can affect transcript abundance via nonsense-mediated decay (NMD) (Holbrook et al., 2004). Stop-gained variants located 50-55 bp or more before the last exon junction often induce NMD, while variants located beyond this threshold are more likely to escape NMD and therefore produce truncated protein (Nagy and Maquat, 1998). However, these predictions are not perfect (Rivas et al. 2015). Variants that trigger or escape NMD in the same gene can manifest in diseases with different symptoms or methods of inheritance(Miller and Pearce, 2014), making it important to validate whether a given variant induces NMD.

CRISPR/Cas9 genome editing technology (Cong et al., 2013; Jinek et al., 2012; Shalem et al., 2014) has provided an avenue with which to introduce specific variants into the genome in order to validate their effects on expression in the native genomic context. However, editing one variant at a time, isolating hundreds of single cell clones, genotyping and expanding clones and measuring transcript abundance is a hugely time-consuming and expensive process. In addition to the resource cost of completing such an experiment, undetected large on-target mutations (Kosicki et al., 2018), off-target mutations and other clone-specific abnormalities can create noise which requires many replicates of each desired genotype. A less labor-intensive genome editing approach analyzes allelic expression in the polyclonal edited cell population, and has been used to validate the effects of specific rare variants (Li et al., 2017) and all possible mutations in a particular exon using saturation mutagenesis (Findlay et al., 2014).

In this study, we decided to develop and apply a similar polyclonal approach for medium-throughput testing of the expression level effects of eQTLs in transcribed regions and rare premature stop variants. We first tested rare premature stop variants with signs of NMD in the Genotype Tissue Expression (GTEx) v8 data to validate the ability of our assay to detect effects on transcript abundance, and then applied the assay to fine-mapped eQTLs from GTEx. Finally, we assayed premature stop variants in two Mendelian disease genes, *GLI3* and *ROR2*, to evaluate our ability to test NMD in a clinically-useful context. Stop-gained variants towards the beginning of *GLI3* are associated with Greig cephalopolysyndactyly, while variants towards the end of the gene are associated with the clinically distinct Pallister-Hall syndrome (Furniss et al., 2007; Johnston et al., 2005). Stop-gained variants towards the beginning of *ROR2* are associated with the autosomal recessive Robinow syndrome, while variants towards the end of the gene are associated with autosomal dominant Brachydactyly type B1 (Ben-Shachar et al., 2009; Schwabe et al., 2000). It has been hypothesized that in both genes the disease manifestation is impacted by whether or not the variant triggers NMD. In such situations, experimental testing of NMD can be valuable for disease diagnosis and prognosis.

## Methods

### fgwas enrichment

First, we sought to establish the relevance of testing eQTL effects driven by variants within transcripts by analyzing the extent of cis-eQTL enrichment in functional elements of the genome. We used GTEx v6 fibroblast eQTL data and a diverse set of annotations: Gene annotations were obtained from GENCODE (Harrow et al., 2012), and regulatory annotations (CTCF-binding site, enhancer, open chromatin region, promoter, promoter-flanking region, and TF binding site) were obtained from the Ensembl regulatory build release 80 (Zerbino et al., 2015). Additional annotations include CADD variant consequence scores (Kircher et al., 2014), SPIDEX machine-learning based prediction of splicing effects (Xiong et al., 2015), experimentally validated miRNA binding sites from Tarbase (Vergoulis et al., 2012), 3’ UTR regulatory elements (Oikonomou et al., 2014), and RNA-binding protein sites from CLIPdb (Yang et al., 2015). Significant fibroblast eQTLs were analyzed for enrichment in these functional annotations using fgwas (Pickrell, 2014), with each annotation tested separately.

### Assay design

The design of the assay is illustrated in Figure 1a. In order to validate transcript regulatory variants’ allelic effects on transcript abundance, we utilized CRISPR/Cas9 genome editing with a gRNA specific to the locus of the variant of interest and a single-stranded DNA (ssDNA) template containing the alternative allele for homology-directed repair (HDR). For each variant of interest, we transfected the gRNA and ssDNA template into a well of inducible Cas9 293T cells. After editing, cells were harvested for gDNA and mRNA, followed by amplicon sequencing of the locus of interest in each. A regulatory effect of the variant is detected as a difference in the ratio of the alternative allele between gDNA and mRNA. This effect size is calculated as the log ratio of the alternative allele in cDNA over the ratio of the alternative allele in gDNA: log2(cDNA alt/ref / gDNA alt/ref), or the allelic fold change (aFC).

**Figure 1.**
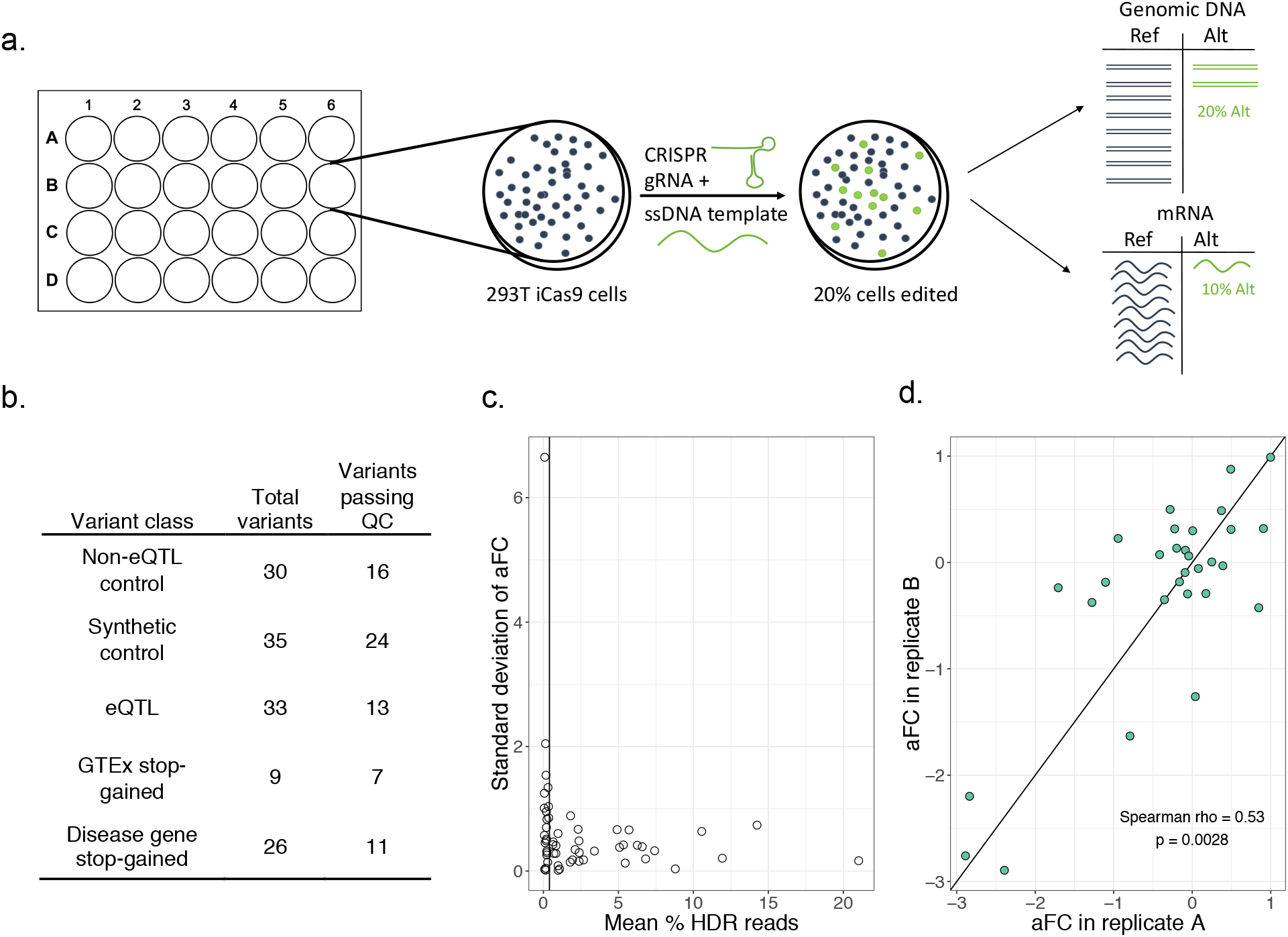
Polyclonal allelic expression assay to detect the effects of regulatory variants. (a) Assay schematic. Inducible-Cas9 293T cells undergo homologous recombination after transfection with the gRNA and ssDNA template in order to introduce the alternative allele to the cells. Editing is followed by targeted sequencing of gDNA and mRNA to detect the ratio of alt/ref alleles in the polyclonal population of cells. (b) Table with the number of each type of control and putative regulatory variant edited with the assay. (c) Homologous recombination rate versus standard deviation for variants replicated 2-3 times with assay. Vertical line shows 0.4% HR cutoff which was used to filter variants for subsequent analysis. (d) Scatter plot showing reproducibility of effect size detected by polyclonal allelic expression assay for two replicate experiments editing the same variants.

### Variant selection

In this study, we edited five types of variants: GTEx stop-gained, GTEx eQTL, disease gene stop-gained, non-eQTL synonymous control, and synthetic control variants (Figure 1b).

Stop-gained variants from the general population were obtained from the GTEx v6 data release. Starting with all stop-gained variants that were singletons in GTEx v6, we used allele-specific expression (ASE) data from the fibroblast sample of the individual carrying the variant to select those that are likely triggering NMD. The selected variants have RNA-seq coverage of >=20 reads, a reference ratio Ref/(Ref+Alt) > 0.7, and are located in a gene with > 5 RPKM in a published HEK293 RNA-seq dataset (Sultan et al., 2014). Additionally, we required ASE data in at least 5 tissues and a first quartile of ASE across tissues of > 0.7 to select variants where NMD does not appear to be highly tissue-specific. Finally, we selected variants > 30 bp from the end of an exon for primer design. Nine variants were used for editing.

eQTL variants were obtained from the GTEx v8 data release. Significant eQTL variants in fibroblasts were filtered for being within at least one protein-coding transcript, having a CAVIAR fine-mapping posterior probability of association > 0.8, an eGene with > 1 RPKM in HEK293 cells, and an effect size in the top quartile of effect sizes of all associations (aFC > 0.30). The top 33 highest effect size variants with successful gRNA and primer design were chosen for editing.

Ten stop-gained variants for each of the disease genes *GLI3* and *ROR2* were created by changing a codon in the transcript to a stop codon. The stop codons were spaced 20 bp apart in both directions from the NMD cutoff point (55 bp upstream of last the exon-exon junction). The 6 disease variants tested were obtained from ClinVar (Landrum et al., 2018), choosing disease-associated variants in the two genes on either side of the NMD threshold.

We selected 30 non-eQTL negative control variants from common synonymous variants in GTEx v8 data with an eQTL association p > 0.1 with the gene in which they reside. The templates for the 35 synthetic control variants were designed by introducing a nucleotide other than the reference or alternative allele at the stop-gained variant locus, which does not create a premature stop codon.

### Cell culture

Genome editing was carried out in a doxycycline-inducible Cas9 293T cell line, transduced with pCW-Cas9 plasmid (Addgene plasmid #50661 (Wang et al., 2014)), courtesy of the Sagi Shapira lab. 293T cells were cultured in OptiMEM (Gibco) supplemented with 5% HyClone Cosmic Calf Serum (Fisher), 1% Glutamax (Gibco), 1% NaPyr (Corning), and 1% penicillin/streptomycin (Corning). The cells were passaged and maintained following standard techniques in 5% CO2 and 95% air.

### Genome editing

The protocol for the polyclonal editing assay can be found at https://doi.org/10.17504/protocols.io.7c6hize. gRNAs were designed with E-CRISP version 5.3 (Heigwer et al., 2014) using medium settings, with an NGG PAM, a 5’ G, excluding designs with more than 5 off-targets, classifying off-targets as having up to 3 mismatches in 5’ region of the gRNA. gRNAs were ordered as gBlocks gene fragments (IDT): a U6 promoter sequence followed by the specific gRNA and tracr sequence (Arbab et al., 2015). The gBlocks were amplified using Q5 high fidelity 2X master mix (NEB) and gBlock amplification primers (Arbab et al., 2015). Homologous templates were designed by extracting the sequence 50 bp upstream and downstream of each variant and substituting the reference allele with the alternative allele. Stop-gained control templates have another nucleotide substituted in the variant position which does not create a stop codon. Homologous templates were synthesized as ultramers by IDT. If possible, primers which amplify both cDNA and gDNA were designed using IDT primer quest, choosing those that cover the PCR target (region spanning the variant and DSB) with at least 15 bp between the PCR target and one primer and at least 60 bp to the other primer. Otherwise, cDNA- and gDNA-specific primers were designed using either the cDNA or gDNA sequence as the template. Nextera adapter sequences were appended to forward and reverse primer sequences as follows:

GTCTCGTGGGCTCGGAGATGTGTATAAGAGACAG+ForwardPrimerSequence TCGTCGGCAGCGTCAGATGTGTATAAGAGACAG+ReversePrimerSequence Primers were ordered as standard oligos from IDT.

Twenty-four hours before transfection for CRISPR editing, iCas9 293T cells were plated in 24-well plates and induced with 5 ug/mL of doxycycline, with a separate well for each targeted variant. Cells were transfected with 500 ng homologous template and 500 ng gRNA gblock using Lipofectamine MessengerMAX transfection reagent. After 24 hours, transfection reagent was removed and replaced with new media. Cells were split after 4 days and 6 days, and DNA and RNA were extracted from the polyclonal edited cultures at 9 days. 75% of the 24-well culture was harvested for RNA using IBI Isolate DNA/RNA Reagent according to the manufacturer’s instructions. Purified RNA was quantified by Nanodrop (Thermo Fisher). cDNA was synthesized with ~200 ng of purified RNA using 1/4 reactions of SuperScript IV VILO Master Mix with EZ DNase (Invitrogen). Another 10% of the cell culture was used for DNA extraction using 15 uL of QuickExtract (Lucigen). For the timecourse optimization experiment, mRNA and gDNA was extracted as above at days 4, 6 and 9.

### Library preparation

Amplicon libraries from cDNA and gDNA were created using either the same nextera primers (if possible) or separate nextera primers for cDNA and gDNA. 1 uL of cDNA or gDNA was amplified using Q5 High Fidelity 2X Master Mix (NEB). An indexing PCR was performed next using Nextera XT index kit primers (Illumina) and NEBNext High-Fidelity 2X PCR Master Mix (NEB) which resulted in dual barcoded amplicons with illumina adapters. cDNA and gDNA libraries were mixed in equal volume and sequenced on the MiSeq using 150 bp paired-end reads. We obtained a median coverage of about 85,000 reads per sample.

### Sequencing analysis

Fastqs generated from Illumina software were trimmed for adapter sequences and quality using trimmomatic. Reads were aligned to the gDNA or cDNA sequence specific for each amplicon and categorized as HDR, no edit, or NHEJ using EdiTyper (Yahi et al. in prep). Variants were eliminated if HDR in gDNA was greater than 30% (suggesting the cell line is in fact heterozygous for the variant). Samples were filtered out if they had fewer than 1000 reads covering the locus of interest. Additionally, samples were filtered out if they had an outlier NHEJ rate of greater than 80%, indicative of an alignment error. The effect size for each variant was calculated as the log2((Alt/Ref in cDNA) / (Alt/Ref in gDNA)), or allelic fold change (aFC). An effect size of zero means the variant has no effect on transcript abundance.

### Statistical analysis

Significance between the control variant distribution and the other experimental variant types was determined using a two-sided Wilcoxon rank sum test. An F test was utilized to detect a difference in variance of aFC between non-eQTL control and synthetic control variants, and eQTL and control variants. For each individual regulatory variant, a p-value was calculated from the z-score of the variant’s effect size based on the mean and standard deviation of the control distribution. The p-values were then bonferroni corrected and variants with a corrected p-value of less than 0.05 were considered significant.

### eQTL effect size in GTEx

For the GTEx effect sizes for the eQTLs, we used the allelic fold change (aFC) estimates from the GTEx v8 data release (Mohammadi et al. 2017, GTEx Consortium 2017, GTEx Consortium in prep.). For eQTL effect size in each GTEx tissue, we used the aFC estimates calculated from eQTL data. For stop-gained variants, we calculated aFC as the log2 ratio of the alternative and reference allele counts in the RNA-seq data in GTEx. To analyze the variation of eQTL variants’ effects across GTEx individuals, we calculated the aFC for each eQTL variant across heterozygous individuals in GTEx as the ratio of allele counts in the gene body (Mohammadi et al., 2017). Samples were filtered for those with greater than 50 reads covering heterozygous sites in the gene.

## Results

First, we assessed the right timepoint to harvest mRNA after transfection with CRISPR constructs. Since mRNA is likely to remain in the cell for hours to days after editing has occurred, we expect to see fewer edited mRNA molecules early after transfection. To find the optimal timepoint, we edited 17 control variants (in 17 different genes) which are not expected to have an effect on expression and harvested at three timepoints post-transfection: 4 days, 6 days and 9 days. At four days, the edited allele is depleted in the mRNA (Supplemental Figure 3a). However, this effect is lessened after 6 days and gone by 9 days. Therefore, we used the 9-day timepoint for the assay in order to analyze only the mRNA which has been transcribed post-editing.

Next, we assessed how technical variation in editing efficiency or PCR amplification may affect the robustness of the assay. We analyzed how the homology-directed repair (HDR) rate affects standard deviation of variant effect size between editing replicates of 62 variants (2 replicates for 23 and 3 replicates for 32 variants). We found that very low HDR is associated with a higher standard deviation between replicates (Figure 1c). Therefore, in subsequent analyses we discarded any variant with an HDR rate of less than 0.4% as determined by the frequency of the alternative allele in the gDNA. The HDR rate varies greatly between loci but is very well correlated between replicates of the same variant (Spearman’s rho = 0.95, p = 7×10^-16^, Supplemental Figure 3b), suggesting that the results of the assay are not strongly influenced by PCR amplification bias or variation in transfection efficiency. Altogether, the effect sizes of the two replicates are well correlated (Spearman’s rho = 0.53, p = 2.8×10^-3^, Figure 1d). These results indicate that the polyclonal allelic expression assay can be used to robustly detect the regulatory effects of genetic variants.

In order to determine the optimal set of negative control variants, we compared the distribution of effect sizes of the synthetic control variants (new variants created in the same genes as the stop-gained variants) and non-eQTL variants (common synonymous variants where eQTL effects were tested in GTEx and not observed). The synthetic control variants have several outlier variants with large effect sizes that are consistent in replicates (Supplemental Figure 3c). This suggests that a subset of synthetic variants affect transcript levels and are thus not ideal negative controls. The non-eQTL control variants, however, have effect sizes consistently close to zero (median aFC = −0.009), demonstrating the utility of population data in selecting nonfunctional negative control variants. The variance of the synthetic controls was significantly greater than the variance of the non-eQTL controls (1.02 versus 0.038; F test p = 3.7×10^-8^). The non-eQTL variants were thus utilized as the control distribution for comparison with the stop-gained and eQTL variants tested with the assay.

In order to analyze the effects of genetic variants on gene expression levels using the polyclonal allelic expression assay, we first analyzed rare stop-gained variants from GTEx that are expected to trigger NMD. As a group, the stop-gained variants show the expected negative effect sizes as compared to the control distribution (Wilcoxon p = 3.2×10^-5^, Figure 2a). Five of these variants individually deviate significantly from the control distribution (Bonferroni-corrected z-test p<0.05). These results demonstrate our ability to capture NMD effects with the assay.

**Figure 2.**
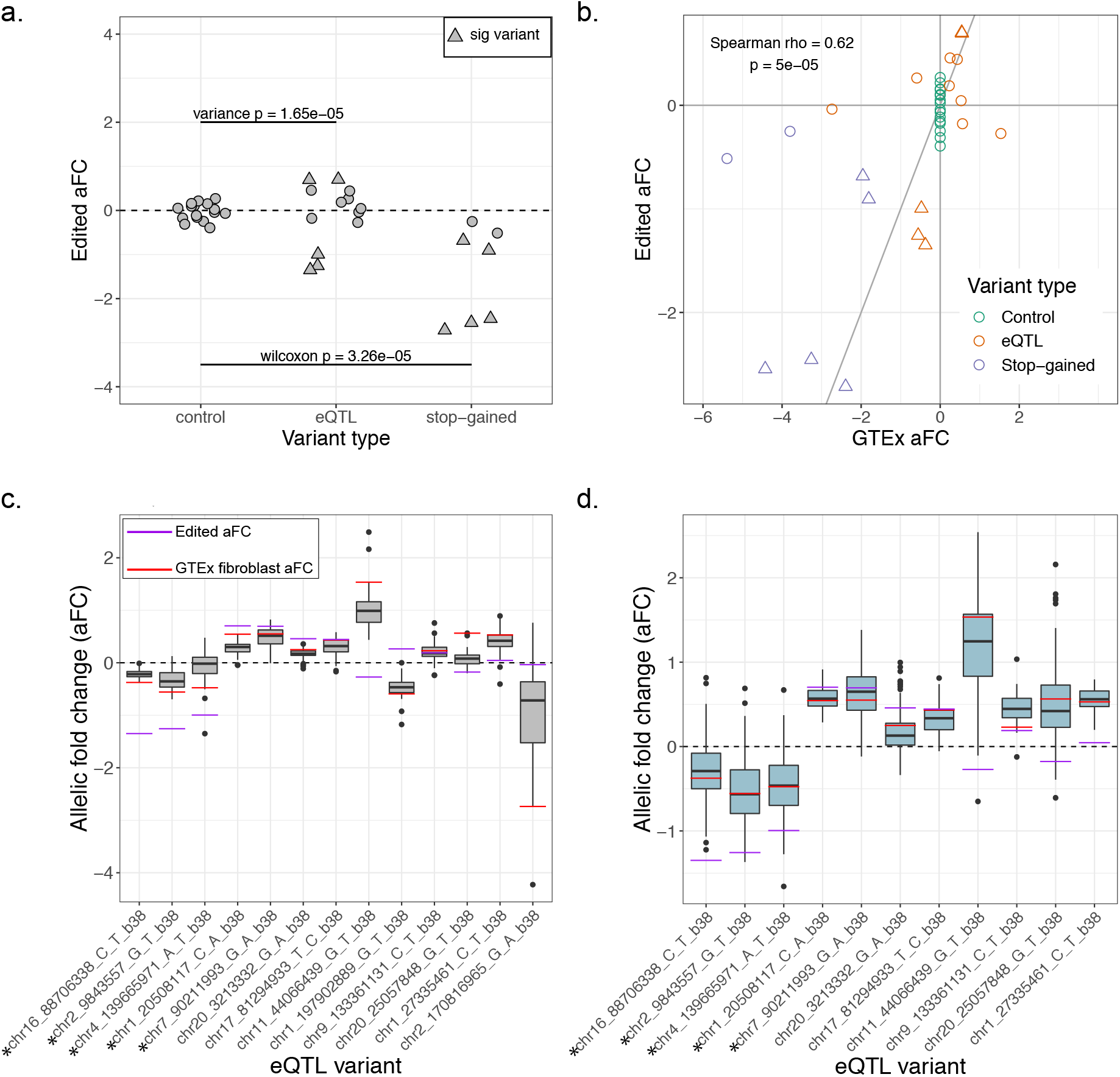
Stop-gained and eQTL variants from GTEx show allele-specific regulatory effects on expression. (a) Effect size of non-eQTL control, eQTL and stop-gained variants after editing with the polyclonal allelic expression assay. Triangular points mark variants whose effect sizes significantly deviate from the control distribution. (b) Correlation between effect sizes of variants in GTEx and effect sizes from the polyclonal allelic expression assay. (c) eQTL effect size (aFC) in GTEx tissues for the 13 edited eQTL variants shown as boxplots, with lines indicating the median effect size in GTEx fibroblasts and in the assay. Asterisks mark variants which were significant in the assay. (d) aFC in GTEx fibroblasts, measured in eQTL heterozygous individuals for 11 of the edited eQTL variants.

Next, we extended the assay to assess putatively causal eQTL variants within transcripts using GTEx fibroblast eQTLs. We chose fibroblasts because GTEx fibroblast transcriptome expression is highly correlated with that of HEK293 cells (rho = 0.68, p < 2.2×10^-16^, Supplemental Figure 2). The 33 eQTL variants chosen for editing are located within the transcript of the eGene with which they are associated and have a high posterior probability of causality based on CAVIAR fine mapping. After editing and QC filtering, 13 eQTL variants remained. The variance of the effect size of the eQTL variants was significantly higher than that of the control variants (0.49 versus 0.038; F-test p = 1.6×10^-5^; Figure 2a), which suggests that the edited eQTL variants as a whole have a greater regulatory effect than the edited control variants. Ten of the 13 variants have an effect in the same direction as the GTEx eQTL effect. Five of the eQTL variants are individually significantly different from the control distribution (Figure 2a), and all five of these variants have an effect in the same direction as in GTEx. Additionally, there is a significant correlation between the effect size of the edited stop-gained, non-eQTL control and eQTL variants and their effects in GTEx (Spearman’s rho = 0.62; p = 5×10^-5^), again indicating that the assay captures regulatory effects seen in the population.

We expect that the lack of effect for some of the eQTL variants is due to the variants not actually being the causal regulatory variants of their association signals. Additionally, our cell line may not perfectly recapitulate genetic regulatory effects of GTEx fibroblast samples. To investigate this, we looked at variation in effect size between GTEx tissues for each of the eQTL variants (figure 2c). We also looked at inter-individual variation within fibroblast samples in GTEx, which may reflect more subtle cell type-specific genetic effects as well as the effects of other regulatory variants that the individuals may have. We measured the effect size in eQTL heterozygotes based on allelic imbalance within the gene body (figure 2d), with eleven of the eQTL variants having sufficient data for this analysis. For all five significant variants, there is agreement in direction between the polyclonal aFC, median heterozygous aFC and eQTL aFC. For several of the other variants, figures 2c and 2d demonstrate a large range of effects both across tissues and across individuals. The observed effect in the cell line, like an individual or tissue, is likely to fall somewhere in a spectrum of possible effects.

In order to apply our assay to the detection of nonsense-mediated decay triggered by disease-associated variants, we introduced stop-gained variants into two disease-associated genes: *ROR2* and *GLI3*. Seven of the edited stop-gained variants are before the 55 bp threshold and were therefore expected to trigger NMD. Of these variants, all seven resulted in negative effect sizes and the distribution of these variants was significantly different from that of both the four variants which were not expected to trigger NMD (Wilcoxon p = 6.1×10^-3^) and the non-eQTL control variants (Wilcoxon p = 5.8×10^-4^, Figure 3a). When tested individually, six of the seven expected NMD variants are significantly different from the control distribution, indicating that we can sensitively detect NMD and NMD-escape across the 55-bp boundary in these two genes.

In addition to the newly created stop-gained variants, we also included disease-causing stop-gained variants from ClinVar. The Arg442Ter mutation in *ROR2* results in a stop-codon right before the predicted NMD cutoff and is associated with the recessively inherited Robinow syndrome. We observe a significant negative effect of this variant (aFC = −1.39, Bonferroni corrected z-test p = 2.2×10^-11^), which is consistent with NMD and the clinical manifestation of disease (Figure 3b). In contrast, the variant Trp749Ter is associated with dominant Type B brachydactyly and falls after the NMD cutoff in the transcript. Our assay shows that Trp749Ter does not affect the expression level of *ROR2* and therefore does not appear to be triggering NMD (aFC = −0.17, corrected p = 1). The one disease variant tested in *GLI3*, Arg792Ter, falling immediately before the predicted border of NMD escape, shows evidence of triggering NMD with a negative effect size in the assay (aFC = −1.02, corrected p = 3.6×10^-6^). This result is consistent with the clinical features of this variant, with Grieg cephalopolysyndactyly syndrome thought to be caused by haploinsufficiency in the gene *GLI3*. The results of editing stop-gained variants in these disease genes indicate that there is a sharp cutoff of NMD / NMD-escape at the previously described 50-55 bp threshold, and pinpoint the immediate molecular mechanism of NMD / NMD-escape for these disease variants. Additionally, the results demonstrate the potential for utilizing this assay to assess whether a variant of clinical interest triggers NMD when it falls close to the threshold of NMD escape.

**Figure 3.**
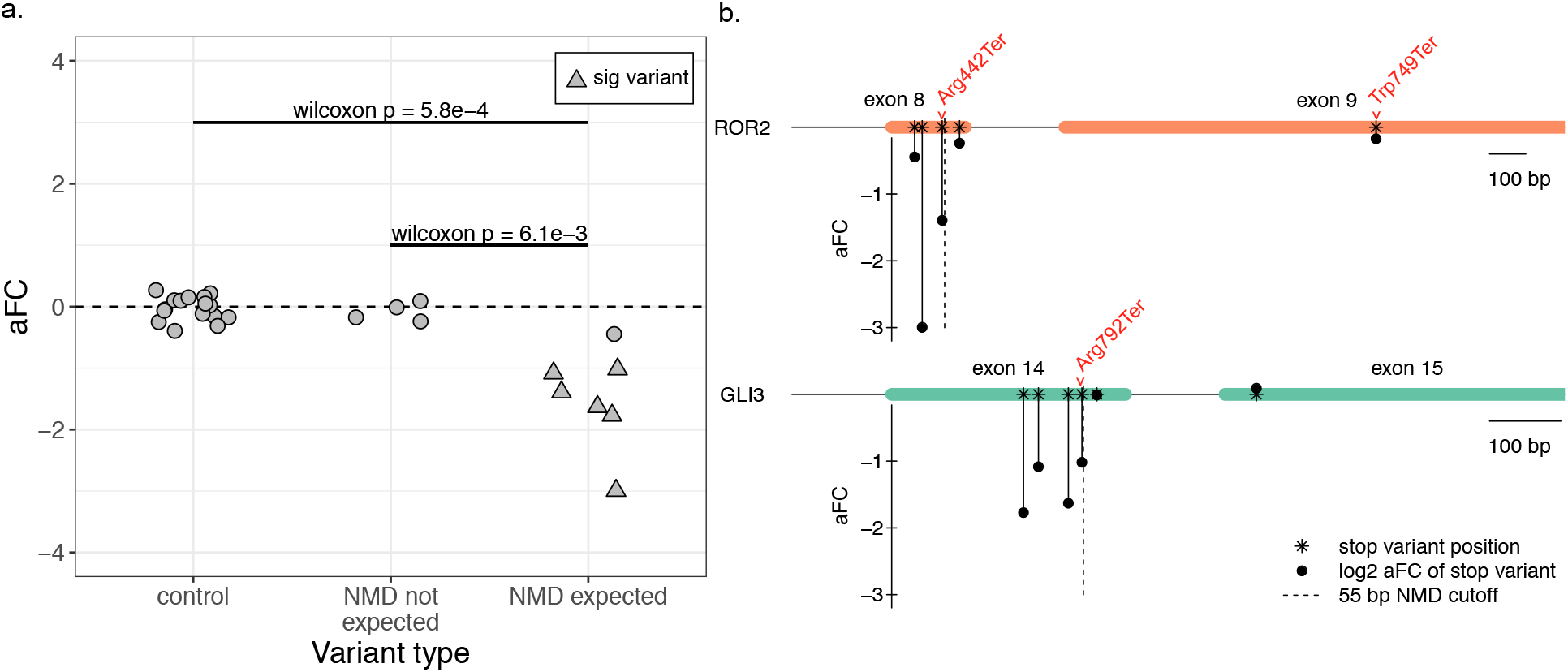
Polyclonal assay effectively detects NMD in disease-associated genes. (a) Effect size in control variants, stop-gained variants after the NMD threshold, and stop-gained variants before the NMD threshold. Triangular points mark variants whose effect size significantly deviates from the control distribution. (b) Diagram of the last two exons of NMD disease genes *ROR2* and *GLI3*, showing the effect size (y-axis) and position in the transcript (x-axis) for each successfully edited variant. Disease-associated variants from ClinVar are labeled in red.

## Discussion

In this study, we described a method utilizing CRISPR/Cas9 genome editing and targeted sequencing to validate regulatory variants without the need for isolating monoclonal cell lines. We demonstrated our ability to reliably detect the effects of stop-gained variants in the general population and in disease cases with the assay. The ability to experimentally assess the effect of potentially disease-causing stop-gained variants could lead to not only better understanding of the rules of NMD / NMD-escape, but also more accurate diagnosis and prognosis. The American College of Medical Genomics recommends caution interpreting pathogenicity of stop-gained or frameshift variants of unknown significance, especially in cases where the variant is in an exon which might be alternatively spliced, or close to the 3’ end of the transcript (Richards et al., 2015). Even though RNA analysis from patients is increasingly used to support variant interpretation (Ben-Shachar et al., 2009; Cummings et al., 2017; Kremer et al., 2017), establishing causality has been difficult since lower expression of a mutant haplotype or gene could be driven by other genetic or environmental factors. Our approach provides evidence that introduction of the specific variant in question underlies transcript level changes, thus reducing the ambiguity of whether variants actually affect transcript abundance. Furthermore, for genes where NMD / NMD-escape is clinically relevant, saturation editing at the 50-55-bp border could build a high-resolution reference for variant interpretation.

This polyclonal assay has the ideal throughput for identifying causal variants from a list of a few to several dozen candidate variants discovered from a rare genetic study. It would be feasible to perform the polyclonal assay on a number of potential regulatory variants, sequencing mRNA and gDNA from the polyclonal culture, and then sort monoclonal cell lines from the same polyclonal culture for only the variants which demonstrate allele-specific regulatory activity. In this approach, the polyclonal assay narrows down the pool of variants to a reasonable number for in-depth follow up with functional assays, protein quantification, etc. The straight-forward nature of the assay makes it easily adoptable in any lab with tissue culture facilities and access to a sequencing instrument.

When we applied the polyclonal assay to eQTL variants, we detected increased effects on expression levels as compared to controls, often in the same direction as the GTEx eQTL effect. Five of 13 variants had significant effects, all consistent with the GTEx eQTL data. This clearly demonstrates the ability of our assay to capture common regulatory variant effects. Some of the non-significant eQTL variants appear to have edited effect sizes consistent with GTEx, but we lack the sensitivity to detect these small effects with confidence. In addition, some of the inconsistencies between the assay results and eQTL data are likely to originate from the eQTL data. Since we do not expect fine mapping to always succeed in identifying the true causal variants at these loci, the undetected effects could represent these situations. Furthermore, with multiple eQTLs for the same gene being common (GTEx Consortium et al., 2017), it is possible that eQTL effect sizes observed in populations reflect multiple regulatory variants in partial LD. Therefore, editing a single variant may not yield the same results as the full haplotype. When we looked at the aFC in heterozygous individuals for these variants in GTEx, we found a broad range of effect sizes, suggesting the presence of effects from multiple variants and potential modifiers that may not be captured by editing a single variant. Finally, assessing genetic regulatory effects even in closely matched cell lines does not necessarily capture effects measured in tissue samples. While this is likely to contribute to some of the differences, cis-eQTLs, especially in the transcribed region, are often highly robust across different tissues (GTEx Consortium et al., 2017), and are expected to replicate in cell lines as well. We highlight that our approach maintains the genomic context of variants and native gene regulation. Thus, it does not suffer from the limitations of massively parallel approaches where discrepancies between eQTL and experimental data may be due to measuring genetic regulatory effects in artificial constructs (Tewhey et al., 2016; van Arensbergen et al., 2019). Altogether, more experimentation and further comparison of population and experimental results are required to fully understand differences between experimental and population data.

Finally, we note that our assay is somewhat limited by HDR efficiency, which varies greatly between loci. Capturing the specific effect of the edited variant requires discarding any reads in the gDNA or cDNA which contain indels created through non-homologous end joining (NHEJ). Since NHEJ often dominates HDR in efficiency, this can result in low numbers of HDR reads. Research in improving the HDR rates in editing is ongoing (Aird et al., 2018; Chu et al., 2015; Maruyama et al., 2015), and likely HDR efficiency will be greatly improved in the future. Additionally, future improvements on base editor technology, which avoids the introduction of double stranded breaks and therefore minimizes the risk of indels (Gaudelli et al., 2017; Komor et al., 2016), could also benefit this system and increase sensitivity of the assay.

## Conclusions

In summary, we have presented a method to validate the allele-specific effects of regulatory variants using CRISPR/Cas9 in a human cell line. When applied to eQTL variants, we see an increased regulatory effect over the control variants, suggesting we are capturing the allele-specific regulatory effects of these variants. Additionally, all of the significant eQTL variants have effects in the same direction as observed in GTEx, demonstrating the assay’s reliability in detecting eQTL effects. The assay is particularly robust in capturing variants triggering NMD across the genome and in rare disease genes, with potential applications in testing the effects of variants of unknown significance from rare variant studies.

## Declarations

### Ethics approval and consent to participate

Not applicable

### Consent for publication

Not applicable

### Availability of data and materials

The sequencing dataset supporting the conclusions of the article is available in the Figshare repository at https://doi.org/10.6084/m9.figshare.9883232.v1. The laboratory protocol is available at https://doi.org/10.17504/protocols.io.7c6hize.

### Authors’ contributions

TL and MB conceived of the idea and wrote the manuscript. MB filtered variants, designed the constructs and analyzed the results. MB, MZ and AG carried out the editing and sequencing experiments. All authors read and approved the final manuscript.

## Acknowledgements

We would like to thank Ana Vasileva for advice and the GTEx Consortium, especially Farhad Hormozdiari and Francois Aguet, for data.

---

**Supplemental Figure 1.**
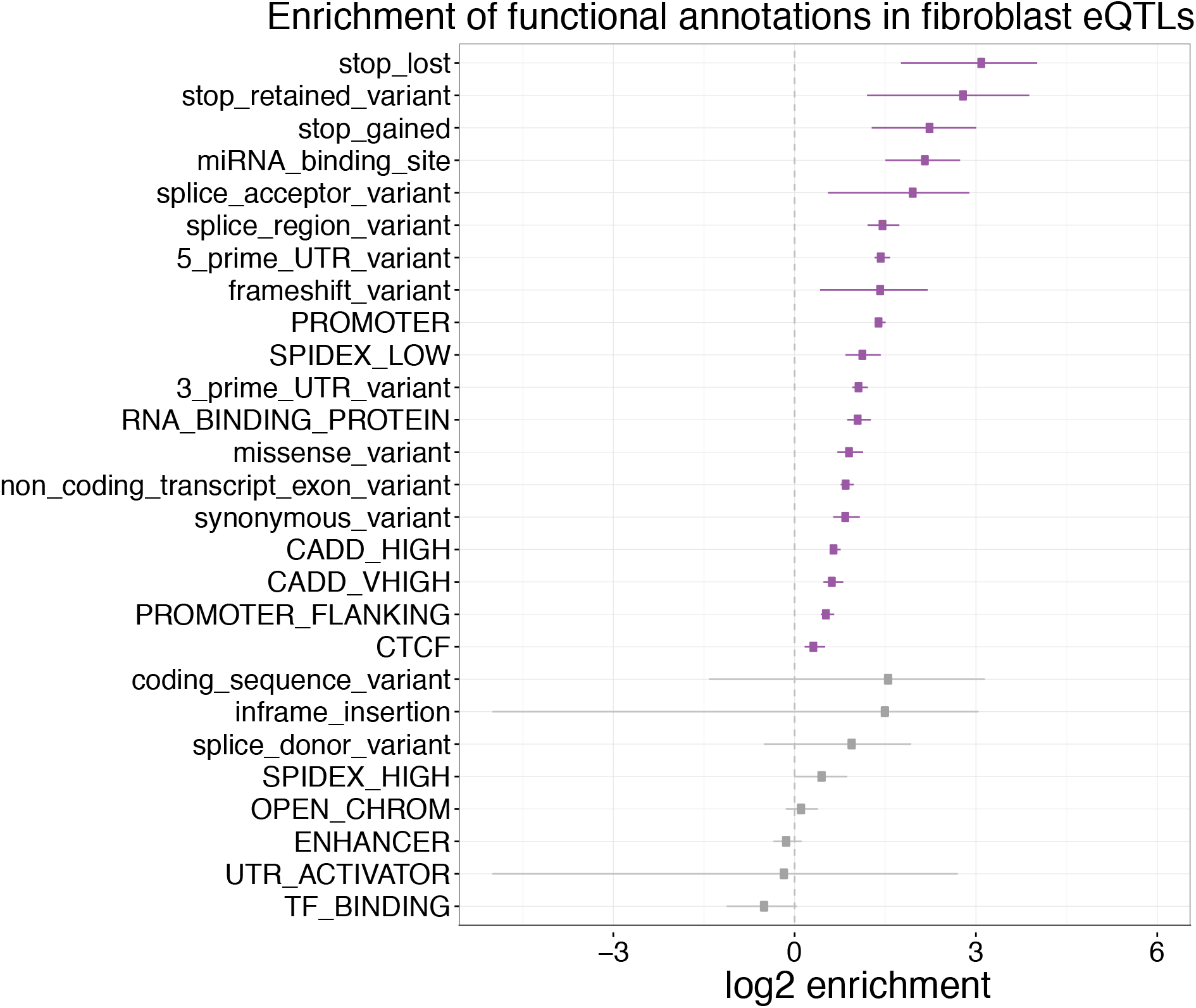
eQTL variants demonstrate enrichment for annotations within the transcript. fgwas enrichment of functional annotations in GTEx fibroblast eQTL variants. Significant annotations are colored in purple.

**Supplemental Figure 2.**
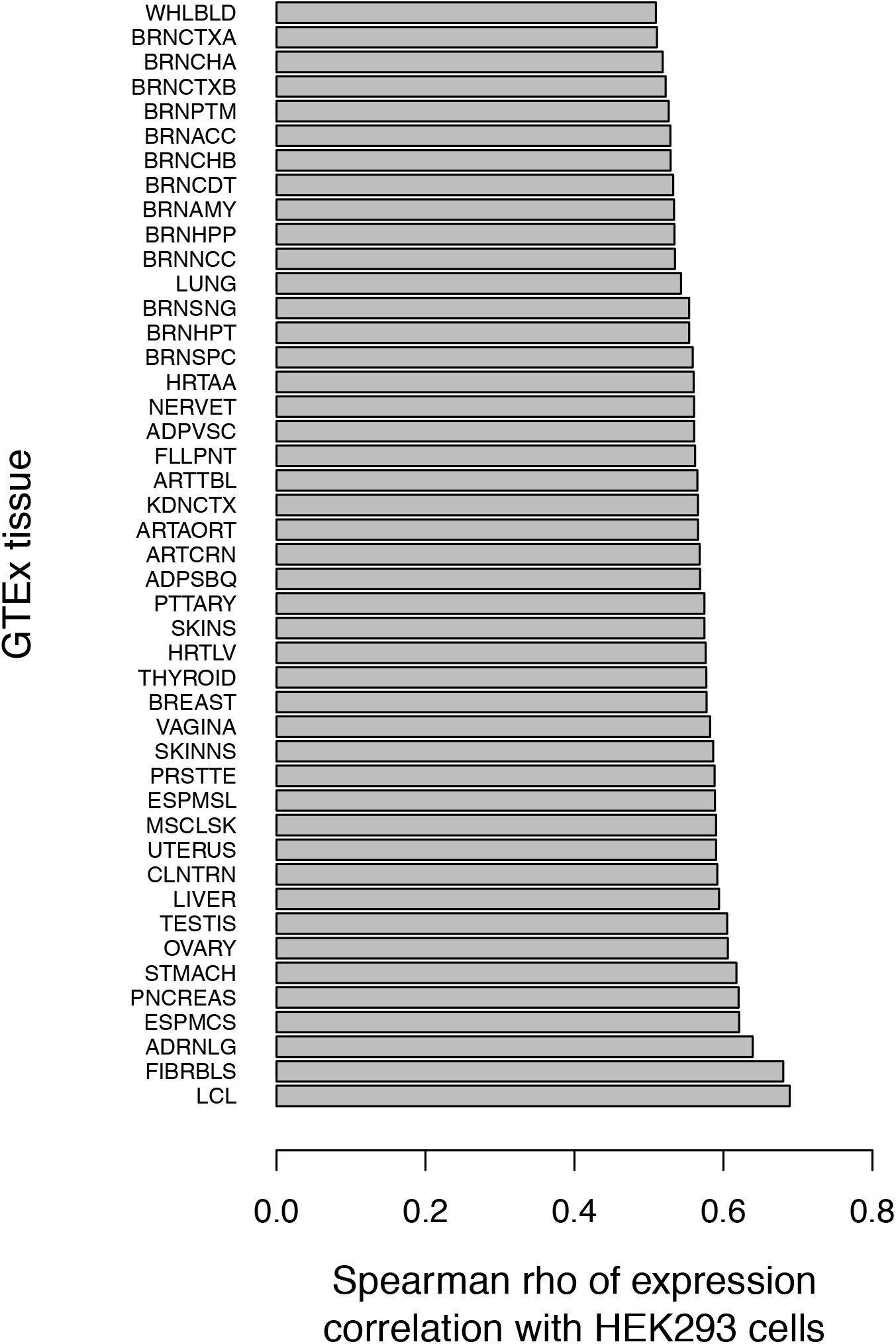
Correlation of transcriptome expression between GTEx tissues and HEK 293 cells.

**Supplemental Figure 3.**
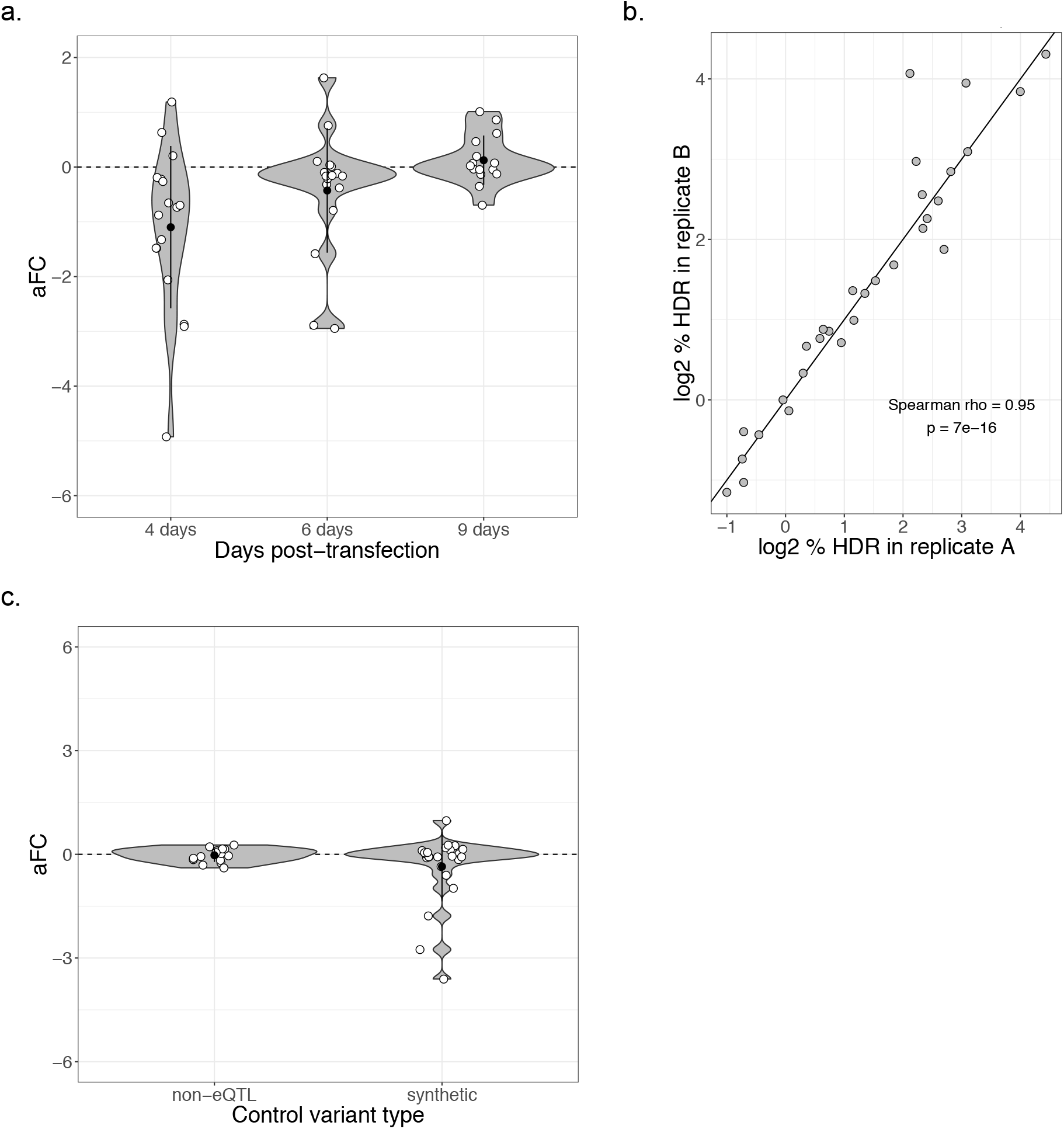
Technical aspects of polyclonal allelic expression assay. (a) Distribution of effect size for control variants over three timepoints post-transfection. (b) Homology direction repair rate in two replicates of control and experimental variants nine days after editing. (c) Distribution of effect size for different control variant types: synthetic control variants and GTEx synonymous non-eQTL control variants.

